# Uptake of plant-derived carbon and proximity to the root determine differences in temporal and spatial stability among microbial groups

**DOI:** 10.1101/2023.03.15.532717

**Authors:** Markus Lange, Mina Azizi-Rad, Georg Dittmann, Dan Frederik Lange, Alice May Orme, Simon Andreas Schroeter, Carsten Simon, Gerd Gleixner

## Abstract

The interactions between plants and soil microorganisms are fundamental for ecosystem functioning. However, it remains unclear if seasonality of plant growth impacts plant-microbial interactions, such as by inducing shifts in the microbial community composition, their biomass, or changes in the microbial uptake of plant-derived carbon. Here, we investigate the stability of microbial biomass of different functional groups and their net assimilation of plant-derived carbon over an entire growing season. Using a C3-C4 vegetation change experiment, and taking advantage of natural abundances of ^13^C, we measured the plant-derived carbon in lipid biomarkers of soil microorganisms in rhizosphere and non-rhizosphere soil. We found that temporal and spatial stability was higher in bacterial than in fungal biomass, while the high temporal stability of all bacterial groups even increased in close proximity to roots. Moreover, differences in the association to plants, i.e., symbionts vs. free-living microorganisms, tend to determine the stability in the uptake of plant-derived carbon. Our results indicate, the inputs of plant-derived carbon over the growing season did not result in a shift in the microbial community composition, but instead, functional groups that are not in obligate symbiosis with plants showed a varying use of soil- and plant-derived carbon.

## Introduction

Terrestrial ecosystems greatly depend on the matter and energy supplied by plants and the respective provision of nutrients by microorganisms (Bardgett and van der Putten, 2014). This interaction between plants and microorganisms controls many ecosystem functions, such as the decomposition and accumulation of organic matter in the soil (Balser and Firestone, 2005; Miltner et al., 2012; Cotrufo et al., 2015; Lange et al., 2015). In the close proximity of the roots, the rhizosphere, this relationship is particularly strong, as this is the interface where plant-derived carbon such as exudates and other rhizodeposits enter the soil. The rhizosphere microbial community has direct access to plant-derived carbon as an easily available energy source, and therefore, it is the zone of the most active soil microbial community (Fierer et al., 2007; Kuzyakov and Blagodatskaya, 2015). The quantity of rhizodeposits and their accessibility for the soil microbial community (Mellado-Vazquez et al., 2016) impacts the microbial activity and thereby microbial carbon transformation and soil carbon turnover (Kuzyakov, 2002; Bird et al., 2011). However, as plants often undergo seasonal changes, such as growth after the cold season and dieback at the end of the growing season, seasonal changes in the quantity and quality of rhizodeposition are likely. Theses phenological patterns of plant carbon allocation are assumed to induce a shift in the general plant-microbial interactions over the course of the growing season (Eisenhauer et al., 2018).

This shift in the soil microbial community is largely attributed to phenological, i.e., temporal changes in the quantity and quality of plant carbon allocation to the soil (Habekost et al., 2008). While the first half of the growing season is presumably dominated by plant inputs of rhizodeposits, the second half is dominated by decomposition of dead plant residues, which are decomposed more slowly than rhizodeposits (Kuzyakov, 2002). However, while temporal shifts in the rhizosphere microbial community composition and their functions have gained increasing attention in recent years (e.g., Chaparro et al., 2014; Shi et al., 2015; Hannula et al., 2019; Nuccio et al., 2020), the contribution of soil- and plant-derived carbon to microbial biomass in relation to plant growth and plant dieback has rarely been considered. Therefore, our knowledge of the seasonal changes in microbial carbon sources, i.e., whether they are derived from soil or plants, is limited, hence soil carbon dynamics cannot be fully understood.soil carbon dynamics cannot be fully understood.

A shift in the microbial community composition can be expected to have subsequent effects on ecosystem functions, such as carbon cycling, (e.g., de Vries et al., 2006; Malik et al., 2016). The dual role of the microbial community on soil organic carbon cycling (Cotrufo et al., 2015; Lange et al., 2015; Liang et al., 2017), i.e., both the decomposition of existing soil organic matter reserves and the transfer of plant-derived carbon to a state that persists in the soil is widely recognized. Shifts in the relation of the microbial community to soil organic matter cycling, namely from consumption to production of organic matter, have been reported (Lange et al., 2021). Yet, a mechanistic understanding of how the soil microbial community is shaped by phenological changes in the quantity and quality of plant carbon allocated to the soil is only recently developing. For instance, Nuccio et al. (2020) reported taxonomical and functional succession of bacteria with substrate specialization depending on root growth, aging and decay. Hannula et al. (2019) nonetheless, assumed that seasonal shifts in bacterial communities do not alter overall soil functioning because the number of species that die out or migrate during the season is small. However, it is very likely that large seasonal variations in microbial biomass, microbial community composition and carbon resource availability will impact soil microbial processes, such as priming (Kuzyakov, 2010), re-cycling (Gleixner et al., 2002) or storage (Lange et al., 2015) of soil organic matter.

Here we take advantage of a C3-C4 vegetation change experiment to estimate the net assimilation of plant-derived carbon into different microbial groups over the course of a growing season. In addition, we examined differences between rhizosphere and non-rhizosphere soils to compare the microbial communities and their net carbon assimilation under high vs. low input of plant-derived carbon via roots and to assess the spatial stability of the entire soil community.

We analyzed microbial communities in soils via phospholipid fatty acids (PLFAs) and neutral lipid fatty acids (NLFAs) and assessed microbial carbon net assimilation by incorporation of ^13^C atoms into these lipid markers. The PLFA-NLFA method is of particular advantage when working with natural isotopic abundances of carbon, as it is highly sensitive to the shifts in the isotopic signatures of lipid biomarkers since fatty acids mainly consist of carbon (Olsson and Johnson, 2005; Yao et al., 2015; Pett-Ridge and Firestone, 2017). The assignment of specific PLFA markers to distinct microbial groups is comparable to the ecological concept of functional groups (Joergensen and Wichern, 2008). This is especially useful when investigating soil carbon dynamics, as the functional groups differ in their needs of carbon availability. The distinct microbial groups in soils are: Gram-positive (G+) bacteria, which are commonly considered as decomposers of soil organic matter (SOM) (Kramer and Gleixner, 2008; Bahn et al., 2013); Gram-negative (G-) bacteria, which are considered to have a high affinity for plant-derived carbon, such as root exudates and therefore preferably colonize the rhizosphere (Denef et al., 2007; Kramer and Gleixner, 2008); saprotrophic fungi, which are able to decompose root exudates and plant litter as well as SOM (Treonis et al., 2004; Garcia-Pausas and Paterson, 2011; Mellado-Vázquez et al., 2019), and AM fungi, which are obligate symbionts that actively form associations with most of the known terrestrial plants (Smith, 2008) and actively trade carbon for nutrients with the host plant (Bonfante and Genre, 2008; Chowdhury et al., 2022). Vegetation change experiments are a common method to study the carbon flow between plants, microorganisms and SOM (e.g. Balesdent and Balabane, 1996; Gleixner, 2013). Differences in the photosynthetic pathways of C3 and C4 plants result in naturally distinct δ^13^C values of both plant types (C3: approx. −25 ‰; C4: approx. −12 ‰; Degens, 1969; O’Leary, 1981). In such experiments, the native plant community of either C3 or C4 plants is exchanged by the respective other plant type, while the soil remains unchanged. Thus, the soil receives the isotopic imprint of the newly established plant type over time. In combination with lipid biomarkers and stable isotope ratio analysis (Evershed et al., 2006; Kindler et al., 2009; Garcia-Pausas and Paterson, 2011), these experiments are used to explore interactions among plants, soil and microbes (Pett-Ridge and Firestone, 2017).

In this study, we asked i) if the microbial communities of the rhizosphere and non-rhizosphere soil differ in their composition and their net carbon assimilation and ii) if phenological changes impact microbial groups differently, depending on their life strategies. We hypothesized that the rhizosphere is dominated by microbial groups with a high affinity for plant-derived carbon and that the biomass of these groups contains higher portions of plant-derived carbon than microorganisms dominating in non-rhizosphere soil. Furthermore, we hypothesized that the proportion of plant-derived carbon in rhizosphere and non-rhizosphere microorganisms is relatively stable over the growing season, but that the biomass of groups with a high affinity for plant-derived carbon would be less stable and thus reflect phenological plant carbon allocation patterns.

## Material and Methods

### Site description and experimental design

The C3-C4 vegetation change experiment at the Max Planck Institute for Biogeochemistry in Jena, Germany was established in 2002. Eight plots of 24 m^2^ were set up directly next to each other to avoid environmental biases, such as climate. Two different types of homogenized soil were filled into two plots each to a depth of 2 m (see Mellado-Vázquez et al., 2019 for details). In short, the first one was originally derived from a forest A-horizon and had a coarse texture (50% sand, 44% silt and 6% clay; pH 6.9) and relatively higher soil organic carbon concentration. The second soil was derived from the B-horizon of a calcareous soil and had finer texture (9% sand, 75% silt and 16% clay; pH 7.8) but lower soil organic carbon concentration. Both soils originally had C3 vegetation cover. Until 2006, the entire experiment was continuously cropped with C3 vegetation (Malik et al., 2012). The selection of plant species was based on comparable biomass production and phenology. At the end of the 2006 growing season, the vegetation change was implemented. The C3 vegetation on one of the two coarse soil plots was replaced with C4 vegetation, and the C3 vegetation on one of the two fine soil plots was replaced with C4 vegetation. The vegetation change was repeated in the same way on a similar set of soils in 2011 in direct proximity to the initial experiment. This resulted in four C3 plots and four C4 plots. All plots were equally managed. Each autumn, the total shoot biomass was harvested and equal amounts were re-distributed on each plot considering the vegetation type. From October to April, i.e. during the non-growing season, plots were covered with a water permeable sheet that allowed plant litter decomposition but prevented weed germination. In the following spring the plots were sown again with their respective vegetation type.

#### Soil sampling

Soil samples were collected during the 2016 growing season, starting in June and continuing at monthly intervals until September, resulting in four sampling dates. Three soil cores (approx. 20 cm by 20 cm) were taken to 10 cm soil depth using a spade from a randomly chosen location within a plot at each sampling time. Immediately after sampling, the soil was manually separated into soil material with direct contact to plant roots (“rhizosphere soil”; see photo SI) and soil material without contact to plant roots (“non-rhizosphere soil”). The corresponding subsamples were mixed to obtain one composite sample each for the rhizosphere and non-rhizosphere soil per plot. Subsequently, samples were immediately sieved (2 mm mesh) and all plant debris was manually removed. Soil samples were then stored at −20 °C for less than a week before lipid extraction.

#### Lipid analysis

Phospholipids and neutral lipids were extracted from soil according to a modified protocol of Bligh and Dyer (1959) in a Büchi SpeedExtractor (BÜCHI Labortechnik GmbH, Essen, Germany) using a mixture (2:1:0.8) of chloroform, methanol and phosphate buffer (0.05 M, pH 7.4). Afterwards, the total lipid extract was separated from water-soluble compounds using phase separation after adding chloroform. The total lipid extract was split into neutral lipids, glycolipids and phospholipids by sequential elution with chloroform, acetone and methanol from a Chromabond^®^ silica column (Macherey Nagel GmbH & Co. KG, Düren, Germany). Phospholipids and neutral lipids were hydrolyzed and methylated using a methanolic KOH solution (0,01 gm^-1^) followed by phase separation after adding chloroform and drying over Na_2_SO_4_. The resulting water-free fatty acid methyl esters (FAMEs) were purified by eluting with a 3:1 mixture of n-hexane and dichloromethane from a Chromabond^®^ aminopropyl column (Macherey Nagel GmbH & Co. KG, Düren, Germany).

FAMEs were quantified using a GC-FID system (GC: HP 6890 Series, AED: G 2350 A, Agilent Technologies Inc., Santa Clara, USA) relative to the internal phospholipid and neutral lipid standard (19:0). Calculation of fatty acid mass was done according to Equation 1:

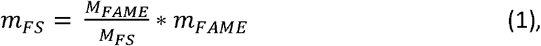

where *m_FA_* is the mass of fatty acids, *m_FAME_* the mass of fatty acid methyl esters, *M_FA_* the molar mass of the fatty acids and *M_FAME_* the molar mass of the fatty acid methyl esters.

Referring the fatty acid mass to the soil dry weight gives the final PLFA and NLFA concentrations for further analysis.

PLFAs and NLFAs were assigned to different microbial groups according to our in-house library (e.g., Thoms and Gleixner, 2013; Mellado-Vazquez et al., 2016; Karlowsky et al., 2018; Mellado-Vázquez et al., 2019). All monounsaturated PLFAs (16:1ω5, 16:1ω7, 17:1, 18:1ω7, 18:1ω9) were assigned to G-bacteria, while all the branched fatty acids (15:0i, 15:0a, 16:0i, 17:0i, 17:0a) were considered as G+ bacterial markers (Zelles et al., 1997; Zelles, 1999). Cyclopropyl fatty acids cy17:0 and cy19:0 represent a subgroup of G-markers, because they are synthesized under environmental stress (Guckert et al., 1986; Kaur et al., 2005) and are often observed in environments richer in G+ bacteria than G-bacteria (Treonis et al., 2004; Mellado-Vazquez et al., 2016). Consequently, they were analyzed separately as “cy G-markers”. In addition, 10Me17:0 and 10Me18:0 were analyzed separately as markers for G+ *Actinobacteria* (Zelles, 1999). The PLFA 18:2ω6,9 was assigned to saprophytic fungi (Zelles, 1999; Frostegård et al., 2011) and the NLFA 16:1ω5 was used as marker for AM fungi (Olsson, 1999). In addition, the PLFA marker 20:4ω6 was used as general marker for soil fauna (Ruess and Chamberlain, 2010).

Incorporation of ^13^C from plant-derived material into the NLFA and PLFA markers for microbial biomass was determined using compound-specific ^13^C analysis with a GC-IRMS system (GC: 7890A, Agilent Technologies Inc., Santa Clara, USA; IRMS: Delta V Plus, Thermo Fisher Scientific, Waltham, USA). Offset during the measurement was corrected using the known δ^13^C-value of the internal phospholipid standards (19:0) and additionally the δ-values were corrected for the methyl group introduced during methylation.

Carbon isotope ratios of individual PLFA markers were measured in triplicate in a GC-IRMS system (HP5890 GC, Agilent Technologies, Palo Alto USA; GC combustion III and IRMS: Delta Plus XL, Finnigan MAT, Bremen, Germany), using an HP Ultra column (60 m, 0.25 mm internal diameter × 0.25 μm film thickness, Agilent Technologies, Palo Alto, USA) and helium as a carrier gas. A SATFA-mix (saturated fatty acid mix, methyl esters) was injected as an external standard before each triplicate sample measurement and fatty acid 19:0 was used for drift correction (Werner and Brand, 2001). The δ^13^C values of SATFA were analyzed with split mode (1:10); whilst δ^13^C values of monounsaturated and polyunsaturated fatty acids were measured with splitless mode. The initial oven temperature of 140 °C was held for 1 min, followed by an increase in temperature at a rate of 2 °C min^-1^ until reaching 260 °C. Followed by a heating rate of 30 °C min^-1^ until a final temperature of 340 °C that was held during 3 min. The software ISODAT NT 2.0 (SP 2.67, Thermo Fisher, USA) was used for data evaluation. Isotope ratios are expressed as δ^13^C value in per mil [‰] relative to the international reference standard Vienna-PeeDee Belemnite (V-PDB) (Eq. 1) using NBS 19 (Werner and Brand, 2001):

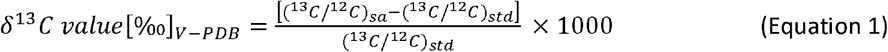

where (^13^C/^12^C)_sa_ is the ^13^C/^12^C ratio of the sample and (^13^C/^12^C)_st_d the ^13^C/^12^C ratio of the reference standard V-PDB. δ^13^C values were also corrected for the methyl carbon added during methylation (Eq. 2; Kramer and Gleixner, 2006):

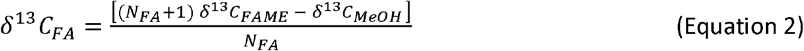

where δ^13^C_FA_ is the isotope ratio of the lipid fatty acid (PLFAs and NLFAs), δ^13^C_FAME_ the isotope ratio of the lipid fatty acid methyl ester, δ^13^C_MeOH_ that of methanol used for derivatization and N_PLFA_ is the number of carbon atoms of the lipid fatty acid.

#### Assessing the carbon origin in PLFA

The net proportion of assimilated carbon derived from plants to individual lipid fatty acids (PLFAs and NLFAs) compared to soil-derived carbon (*Fp*_FA_; Eq. 3) was calculated according to Kramer and Gleixner (2006):

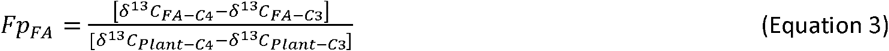

where Δ^13^C_FA-C4_ and δ^13^C_FA-C3_ represent the isotopic values of individual lipid fatty acid markers and the δ^13^C-_Pant-C4_ and δ^13^C-_Plant-C3_ are the isotopic values of plant material from the experiment plot with C4 vegetation change and control C3 plot, respectively.

The δ^13^C values of root material (δ^13^C-_Plant_) was determined on 3 mg of air-dried (40 °C) and ball-milled root material using EA-IRMS. The system was calibrated versus V-PDB using CO_2_ as the reference gas (Werner and Brand, 2001). In addition, soil water content [%] was measured gravimetrically according to the method introduced by Black (1965) from 5 g of soil (wet weight) that were collected as subsamples from the soil samples taken for lipid analysis.

#### Statistical Analysis

To analyze how the concentrations and the net assimilation of plant-derived carbon into lipid markers was impacted by roots and how both metrics would vary over the vegetation period, we employed Linear Mixed-Effects Models (LMM) using the “lmer”-function in the R (R Development Core Team, 2021) library “lme4” (Bates et al., 2015). Starting from a constant null model with “plot identity” as random intercept, we extended the null model stepwise. Since more than one lipid marker is indicative for each of the bacterial subgroups, the random intercept for bacterial models was “lipid marker” fitted within “plot identity”. The fitting sequence of the fixed effects started with “rhizosphere” followed by “season” (= time point of sampling; June, July, August, September) and “soil texture” (coarse, fine). Next, “plant type” (C3, C4) was included in the models to test the effects on lipid marker concentration. To test whether the effects of season were different in rhizosphere vs. non-rhizosphere soil, the interaction term “rhizosphere × season” was included in the models additionally. The maximum likelihood method (ML) was applied and likelihood ratio tests (χ^2^ ratio) were used to compare succeeding models and test for a significant model improvement by the added fixed effects (Zuur et al., 2009). To test if the composition of the lipid markers, based on their relative abundances, showed a shift over the season or was impacted by the environmental factors, a Permutational Multivariate Analysis of Variance (PERMANOVA) was performed applying the “Adonis”-function (permutations = 499) in the “vegan” package (Oksanen et al., 2020). The inverse coefficient of variation (CV^-1^ = mean/ standard deviation (S.D.)) of the microbial biomass and of the microbial net assimilation of plant-derived carbon over the season was calculated to assess the temporal stability (Haddad et al., 2011; Strecker et al., 2016). Differences of the temporal stability among microbial groups and the effect of the proximity to the root was analyzed by applying LMM, similar as described above, i.e., starting from a constant null model, with lipid marker identity and with plot identity as random intercept, and stepwise extension of the null model afterwards. The fitting sequence of the fixed effects started with “microbial group” followed by “rhizosphere” and the interaction term “groups × rhizosphere”. However, in the full model, the dispersion was very high for some groups (see Fig. 3) so the interaction term “groups × rhizosphere” was not significant. Therefore, the rhizosphere effect was tested separately for all groups. These results were finally reported in Fig. 3, as they more appropriately represent the differences shown.

**Fig. 1:**
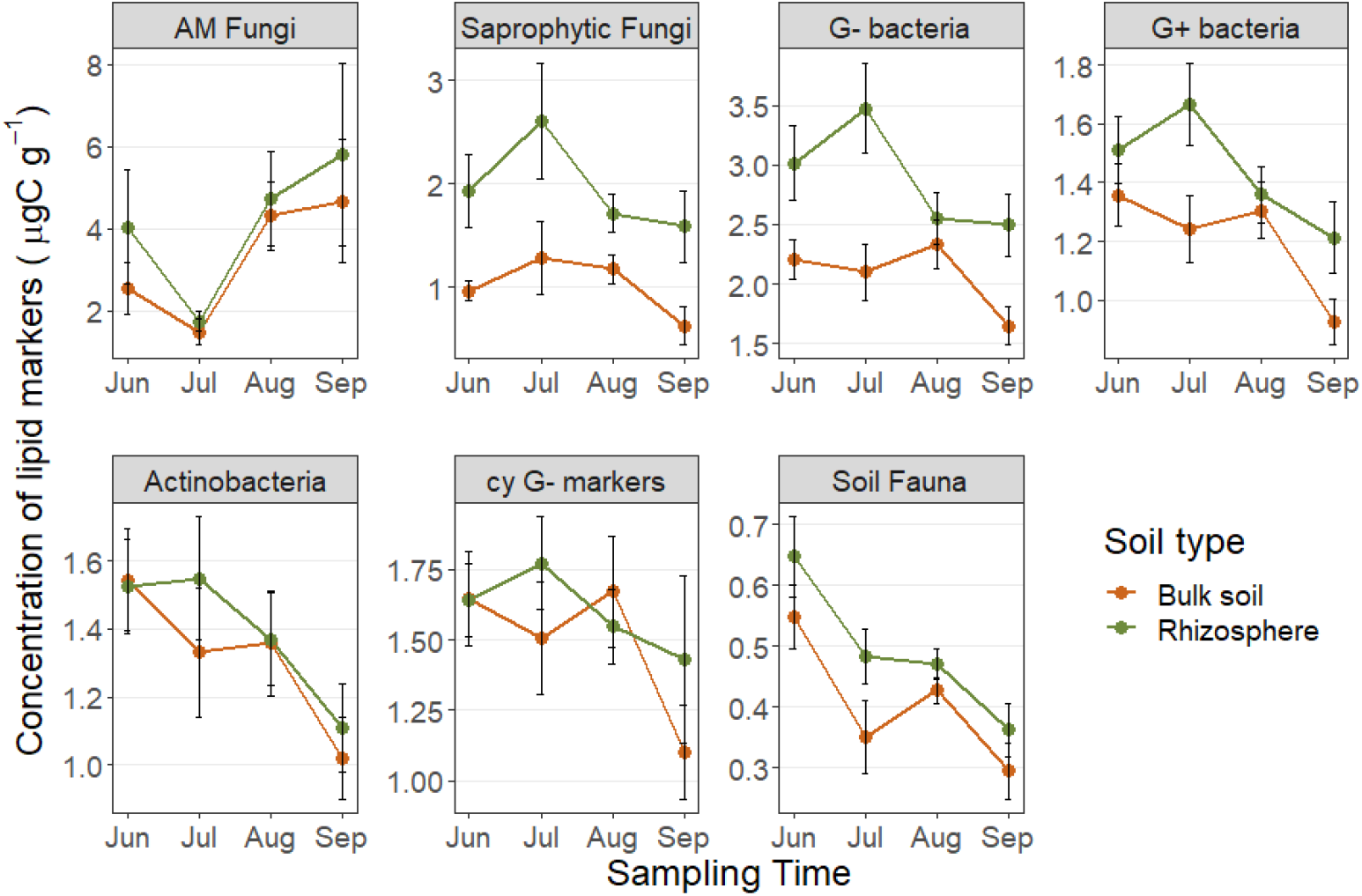
Average concentrations of lipid markers (μg carbon per g soil) assigned to AM fungi and saprophytic fungi, bacteria (G-bacteria, G+ bacteria, Actinobacteria, cyclopropyl markers of G-bacteria) and soil fauna over the growing season in rhizosphere and non-rhizosphere soil. Error bars represent the standard error (S.E.). Note the different scaling among sub-plots.

**Fig. 2:**
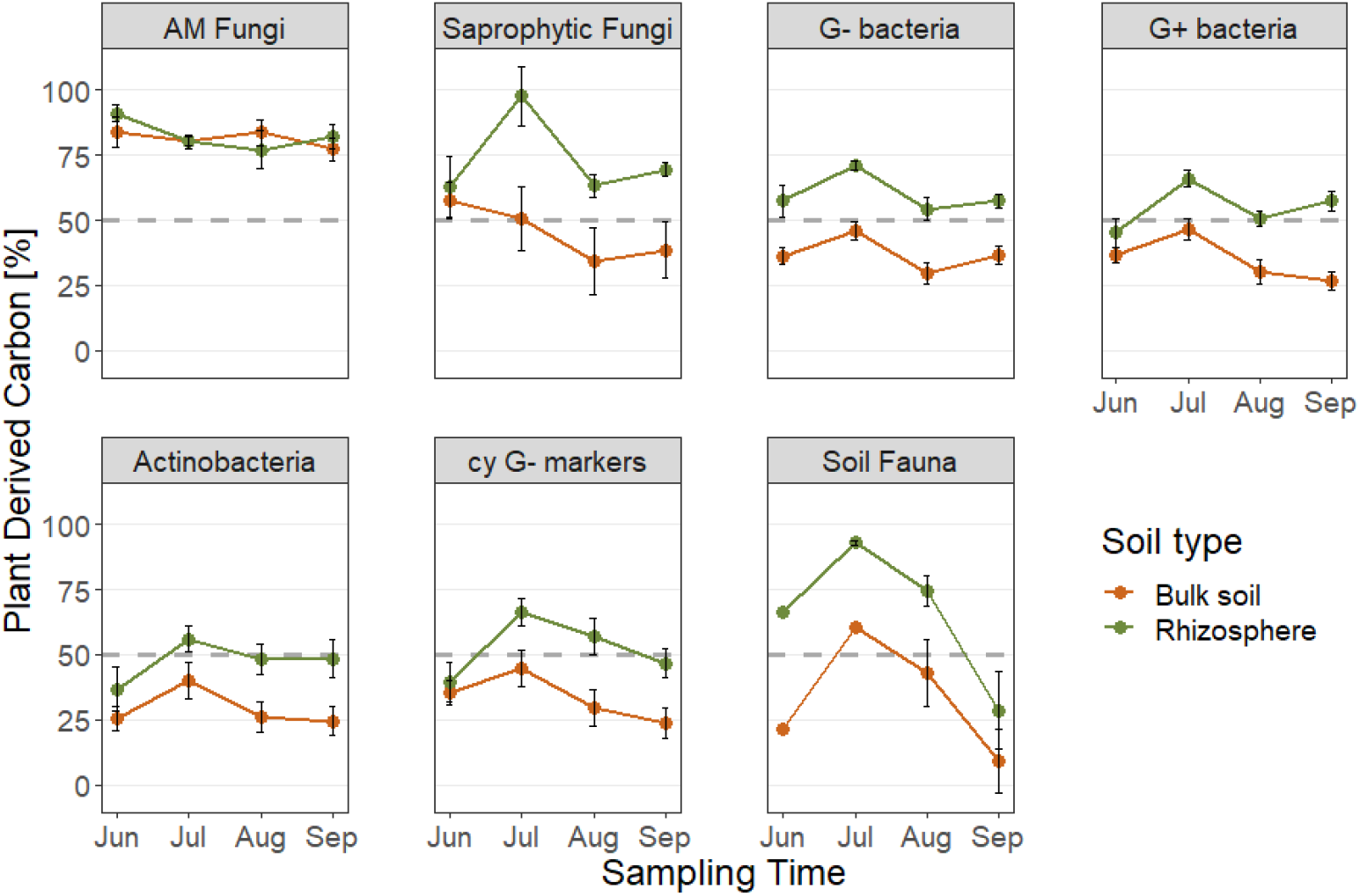
Weighted average of Fp value (plant-derived carbon weighted on concentration of the lipid marker) of AM fungi and saprophytic fungi, bacteria (G-bacteria, G+ bacteria, Actinobacteria, cyclopropyl markers of G-bacteria) and soil fauna over the growing season in rhizosphere and non-rhizosphere soil. Error bars represent the S.E. The gray dashed line provides a guide at 50%. See Supplementary Fig. S2 for the net assimilation of plant-derived carbon in the microbial biomass in μgC g^-1^.

**Fig. 3:**
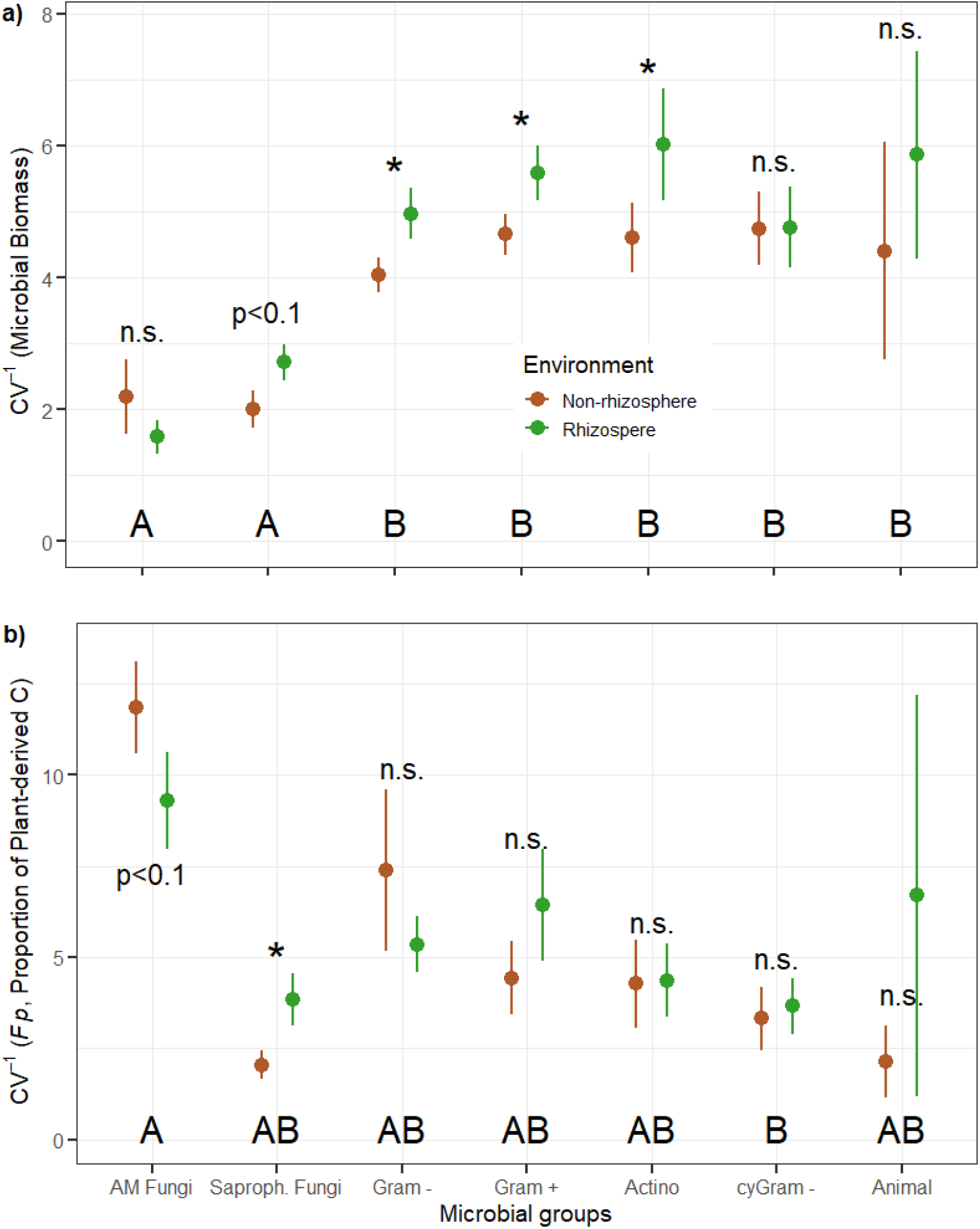
Temporal stability (inverse coefficient of variation, CV^-1^) of a) the microbial biomass and b) the proportion of plant-derived carbon in lipid markers over the growing season in rhizosphere and non-rhizosphere soil. Given are the average values of stability and the error bars represent the standard error (S.E.). See Supplementary Table S3 for numbers on which this graph is based.

## Results

### Microbial biomass and community composition over the growing season

The concentrations of lipid biomarkers differed strongly among microbial groups. Across all samples, the average concentrations of the AM fungi biomarker (NLFA 16:1ω5) was highest with 3.7 μgC (± 3.5 S.D.) followed by the G-bacteria (2.5 ± 1.7 μgC g^-1^) and saprophytic fungi (1.5 ± 1.0 μgC g^-1^). Markers of G+ bacteria had an average concentration of 1.3 ± 0.7 μgC g^-1^, for Actinobacteria it was 1.4 ± 0.6 μgC g^-1^ and for cy G-markers it was 1.5 ± 0.8 μgC g^-1^. Besides the average concentration of the lipid markers, their temporal patterns differed as well. While the concentration for AM fungi was lowest in July and increased sharply towards the end of the growing season, the marker concentrations of saprophytic fungi and most bacterial markers were highest in early summer (July) and decreased towards the end of the growing season. This pattern was particularly pronounced in the rhizosphere (Fig. 1). These seasonal effects were not clearly related to soil moisture, which slightly increased from June (12% soil water content) to July (15%) and decreased again towards August (9%) and September (11%). Including soil moisture into the statistical models did not, or at most slightly reduced, the explained variation by season (Table S1), most likely because the concentration of individual lipid markers was inconsistently correlated to changes in soil moisture (Table S2).

The average concentration of all markers was higher in rhizosphere soil than in non-rhizosphere soil (Table 1, S2). However, this difference was only significant for the markers assigned to saprophytic fungi (concentration increased by 48% from non-rhizosphere to rhizosphere soil), G-bacteria (27% higher concentration in rhizosphere soil than in non-rhizosphere soil), and G+ bacteria (16% higher, Table 1). Although overall significantly different, the marker concentrations of G-bacteria and G+ bacteria did not differ in August (Fig. 1) as indicated by the significant interaction term “rhizosphere × season” (Table 1). Soil texture had an impact on most of the microbial groups except cy G-markers. In contrast, plant type (photosynthetic pathways C3 vs. C4) had no impact on most of the microbial groups, except on cy G-markers. The lipid marker assigned to soil fauna was strongly affected by the season: It constantly decreased from spring towards the end of the growing season; and its concentration was higher in rhizosphere vs. non-rhizosphere soil. Furthermore, in groups to which multiple lipid markers were assigned, the largest effect was the factor “marker” itself, showing that the concentrations of the individual markers greatly differed from one another, even if they were assigned to the same group (Table S2). The composition of the lipid markers was mostly explained by differences in soil, followed by season and the distance to the roots (Table 1). This indicates that effects of soil texture, season and distance to the roots differently impacted the individual organism groups.

**Table 1:** Results of linear effects models testing the effect of the proximity to the roots (“Rhizosphere”) the growing season (“Season”), differences in C3 and C4 vegetation (“Plant type”), soil texture (“Soil texture”) and the interaction between proximity to the roots and the growing season (“Rhizo × Season”) on the concentrations of lipid markers assigned to different microbial groups. Asterisks indicate the level of significance (“***” < 0.001, “**” < 0.01, “*” < 0.05, “.” < 0.1).

|  | Rhizosphere |  | Season |  | Plant type |  | Soil texture |  | Rhizo × Season |  |
| --- | --- | --- | --- | --- | --- | --- | --- | --- | --- | --- |
|  | Chi<sup>2</sup> | R<sup>2</sup> | Chi<sup>2</sup> | R<sup>2</sup> | Chi<sup>2</sup> | R<sup>2</sup> | Chi<sup>2</sup> | R<sup>2</sup> | Chi<sup>2</sup> | R<sup>2</sup> |
| Actinobacteria | 1.5 | 0.00 | 39.8 *** | 0.08 | 0.3 | 0.01 | 1.6 | 0.07 | 3.3 | 0.01 |
| G+ bacteria | 33.3 *** | 0.03 | 72.0 *** | 0.05 | 1.0 | 0.02 | 0.8 | 0.01 | 17.8 *** | 0.01 |
| G- bacteria | 51.2 *** | 0.06 | 24.8 *** | 0.03 | 0.6 | 0.01 | 8.1 ** | 0.12 | 16.0 ** | 0.02 |
| cy G- markers | 1.6 | 0.01 | 12.9 ** | 0.04 | 1.0 | 0.04 | 0.3 | 0.01 | 4.9 | 0.02 |
| AM Fungi | 0.9 | 0.01 | 11.5 ** | 0.16 | 0.4 | 0.01 | 5.2 * | 0.07 | 0.5 | 0.01 |
| Saproph. Fungi | 18.2 *** | 0.22 | 9.3 * | 0.09 | 0.8 | 0.02 | 7.0 ** | 0.11 | 2.2 | 0.02 |
| Soil Fauna | 4.5 * | 0.07 | 30.1 *** | 0.35 | 4.6 * | 0.04 | 2.1 | 0.02 | 1.3 | 0.01 |
| Community Composition | 8.7 *** | 0.09 | 4.1 ** | 0.13 | 2.4 | 0.02 | 14.6 *** | 0.15 | 0.7 n.s. | 0.02 |

### Microbial carbon assimilation over the growing season

Similar to the concentrations of the lipid biomarkers, the content of plant-derived carbon of the lipid biomarkers varied among microbial groups. It was on average highest in the NLFA marker assigned to AM fungi (0.81 ± 0.09), followed by saprophytic fungi (0.59 ± 0.26). The portion of plant-derived carbon in markers of bacterial groups was lower but in a similar range, being highest in G-bacteria (0.47 ± 0.21) and lowest in Actinobacteria (0.37 ±0.20). The plant-derived carbon in the marker for soil fauna was on average 0.43 ± 0.34. The net assimilation of plant-derived carbon depended strongly on the proximity to the roots and was expectedly greater in the rhizosphere (Fig. 2). The net assimilation of plant-derived carbon was only independent of the proximity to the roots in the AM fungi marker (Table 2). The seasonal patterns among all groups were slightly different, however, they all had the greatest contents of plant-derived carbon in early summer (July; Fig. 2).

**Table 2:**
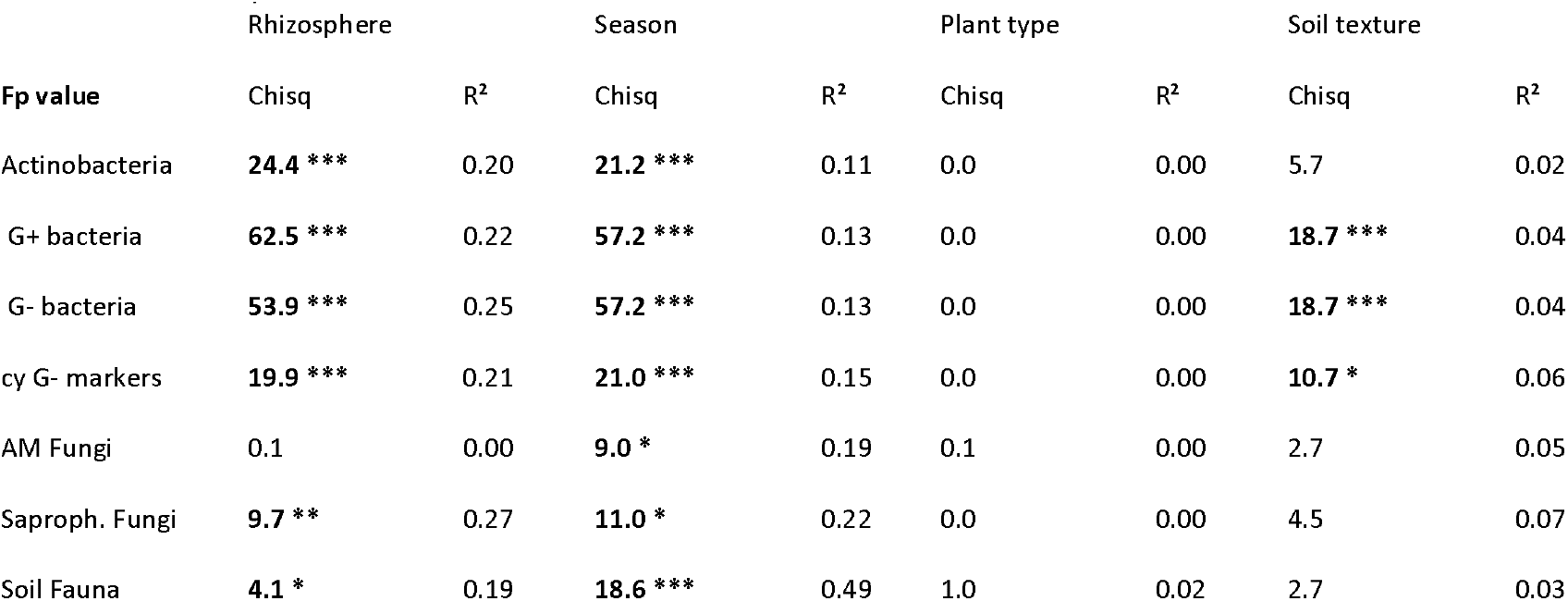
Results of linear effects models testing the effect of the proximity to the roots (“Rhizosphere”) the growing season (“Season”), differences in C3 and C4 vegetation (“Plant type”), soil texture (“Soil texture”) and the interaction between proximity to the roots and the growing season (“Rhizo × Season”) on the proportion of plant-derived carbon (Fp) in lipid markers assigned to different microbial groups. Asterisks indicate the level of significance (“***” < 0.001, “**” < 0.01, “*” <0.05, “.”<0.1).

|  | Rhizosphere |  | Season |  | Plant type |  | Soil texture |  |
| --- | --- | --- | --- | --- | --- | --- | --- | --- |
| Fp value | Chisq | R<sup>2</sup> | Chisq | R<sup>2</sup> | Chisq | R<sup>2</sup> | Chisq | R<sup>2</sup> |
| Actinobacteria | **24.4 \*\*\*** | 0.20 | **21.2 \*\*\*** | 0.11 | 0.0 | 0.00 | 5.7 | 0.02 |
| G+ bacteria | **62.5 \*\*\*** | 0.22 | **57.2 \*\*\*** | 0.13 | 0.0 | 0.00 | **18.7 \*\*\*** | 0.04 |
| G- bacteria | **53.9 \*\*\*** | 0.25 | **57.2 \*\*\*** | 0.13 | 0.0 | 0.00 | **18.7 \*\*\*** | 0.04 |
| cy G- markers | **19.9 \*\*\*** | 0.21 | **21.0 \*\*\*** | 0.15 | 0.0 | 0.00 | **10.7 \*** | 0.06 |
| AM Fungi | 0.1 | 0.00 | **9.0 \*** | 0.19 | 0.1 | 0.00 | 2.7 | 0.05 |
| Saproph. Fungi | **9.7 \*\*** | 0.27 | **11.0 \*** | 0.22 | 0.0 | 0.00 | 4.5 | 0.07 |
| Soil Fauna | **4.1 \*** | 0.19 | **18.6 \*\*\*** | 0.49 | 1.0 | 0.02 | 2.7 | 0.03 |

The net assimilation based on the soil faunal marker showed a similar pattern to the microbial markers described above. However, this marker showed the greatest variation in net assimilation of plant-derived carbon during the season (e.g., in the rhizosphere 90% in July and 30% in September). Furthermore, and contrary to the concentrations of the lipid markers, soil texture had no effect on any of the microbial groups, indicating that the net assimilation of plant-derived carbon into lipid markers of the same groups were similar.

### Stability of microbial biomass and assimilation of plant-derived carbon over the growing season

The temporal stability, calculated as the inverse coefficient of variation (see Methods), of the biomass differed among microbial groups. It was significantly lower in AM fungi and saprophytic fungi than in all other bacterial groups and the soil fauna (Fig. 3a). Furthermore, the concentration of markers of G-bacteria, G+ bacteria and Actinobacteria showed a higher temporal stability in the rhizosphere than in non-rhizosphere soil. This pattern was similar in saprophytic fungi by trend (p < 0.01), while the temporal stability of the AM fungi marker, the cy G-markers and the markers assigned to soil fauna was not affected by the proximity to the roots (Fig. 3a).

In contrast to the stability of biomass over the season, the proportion of plant-derived carbon was most stable in AM fungi markers and least stable in cy G-markers (Fig. 3b). The stability of assimilation of plant-derived carbon in other groups was lower but not significantly different from AM fungi markers. The temporal stability of plant-derived carbon assimilation did not differ between rhizosphere and non-rhizosphere soil for the faunal and all bacterial markers. In contrast, while the stability of the net assimilation of plant-derived carbon in saprophytic fungi was on a very low level, the stability in the non-rhizosphere soil was even significantly lower (Fig. 3b). In the AM fungi, there was a trend (p < 0.1) that the temporal stability of the plant-derived carbon net assimilation was higher in the non-rhizosphere than in the rhizosphere.

## Discussion

In this study, we used a C3-C4 vegetation change experiment and induced δ^13^C changes to investigate how the biomass of different microbial groups and their net assimilation of plant-derived carbon change over the season, as affected by root proximity.

### Rhizosphere effects

In line with our first hypothesis, we found that microbial groups with a high affinity for plant-derived carbon, i.e., G+ bacteria and saprophytic fungi were most abundant in the rhizosphere and their biomass was more strongly influenced by proximity to roots than the other microbial groups. The strongest rhizosphere effect was observed in the fungal marker, while the differences between G- and G+ bacteria were not as strong as might have been expected based on their supposed preference for plant or soil-derived carbon. (Denef et al., 2007; Kramer and Gleixner, 2008; Bahn et al., 2013). As such, in recent years a saprotrophic activity of G-bacteria was reported (Bai et al., 2016; Mellado-Vázquez et al., 2019), while the G+ bacteria assimilate large portions of new photosynthates, if available (Mellado-Vazquez et al., 2016; Karlowsky et al., 2018; Mellado-Vázquez et al., 2019; Mielke et al., 2022).

The higher uptake of plant-derived carbon in the rhizosphere by most of the microbial groups was expected because of the higher availability of plant-derived carbon (Bais et al., 2006; Philippot et al., 2013; Pausch and Kuzyakov, 2018). Greater quantities of easily decomposable energy sources (rhizodeposits) lead to higher microbial activity and growth in the rhizosphere compared to non-rhizosphere soil (Kuzyakov and Blagodatskaya, 2015). This was clearly confirmed by the general differences in microbial biomass. However, some groups showed no clear response in biomass between rhizosphere and non-rhizosphere soil. For example, although very different in their life strategies, the biomass of AM fungi and actinobacteria were not affected by the proximity to the roots. AM fungi demonstrated their obligatory dependence on plant-derived carbon (Bago et al., 2000; Chowdhury et al., 2022) since the proportion of plant-derived carbon in the specific lipid marker was not affected by proximity to roots. In contrast, non-impacted biomass of actinobacteria likely indicates a slow growth, which is independent of the availability of easily decomposable plant-derived carbon sources and confirms a general, oligotrophic lifestyle of these organisms (Liu et al., 2018).

Our data show that most microbial groups utilize both plant-as well as soil-derived carbon, and are flexible in choosing these sources though they have different preferences for carbon sources. All groups considered here exhibited an opportunistic carbon use as shown by the significantly higher plant-derived carbon proportion in close proximity to the roots, except for AM fungi which also incorporated high quantities of plant-derived carbon in non-rhizosphere soil. In our study, this opportunistic carbon use is possible because no substrate limitation appears to occur in either rhizosphere or non-rhizosphere soil as indicated by equally low concentrations of the cy G-markers, indicative of stress conditions such as substrate-limitation (Guckert et al., 1986; Kaur et al., 2005).

### Changes over the growing season

While the proximity to roots explained the largest differences in net assimilation of plant-derived carbon, differences in microbial biomass were largely explained by the time point of sampling. These temporal effects on the microbial biomass, however, were not or only weakly related to changes in soil moisture (Suppl. Table S1). Instead, it is likely that the temporal patterns are directly related to root exudation, which in turn might reflect plant requirements driven by environmental conditions such as water or nutrient availability (Maurer et al., 2021). For example, the sharp decrease in AMF biomass in July, when the biomass of most other free-living microorganisms peaked, suggests altered root exudation patterns. Most notably, saprophytic fungal biomass and the proportion of plant-derived carbon in that group increased in the rhizosphere in July. Similar to our study, an opposite pattern in the net assimilation of plant-derived carbon between AM fungi and saprophytic fungi has been reported (Chowdhury et al., 2022). This strongly indicates a directed root exudation, which might be related to environmental conditions such as plant water availability. The effect of soil water content on microbial biomass might therefore also be indirectly mediated through plants via root exudation.

The net assimilation of plant-derived carbon of the soil fauna was very variable throughout the growing season. This could indicate either 1) that with time, different animals of different feeding guilds are reflected by the C20;4ω6 marker or 2) that the soil fauna changed their feeding habit during the season. As this marker is unspecific for soil fauna (Ruess and Chamberlain, 2010) and showed relatively large concentration shifts during the vegetation period, it is more likely that variation in plant-derived carbon in the soil fauna marker reflects a community shift of the soil fauna from fungivore or herbivores in early summer to bacterivores and humus-feeding fauna later.

Contrary to our second hypothesis, there was no clear pattern that the microbial biomass of groups with a high affinity for plant-derived carbon was less stable over the growing season compared to groups with a preference for soil carbon. In contrast, the stability of the biomass of both AM fungi and saprophytic fungi was significantly lower than the biomass stability of all bacterial groups. This is primarily due to the sharp increase and decrease, respectively, in both groups during the July sampling (as discussed above) and likely reflects differences in the life strategies and growth forms between fungi and bacteria (Ho et al., 2017). However, the higher temporal stability of bacterial biomass in the rhizosphere (all groups except cy G-markers) compared to the non-rhizosphere soil was unexpected, as we considered an entire growing season from the early development of the plants to their senescence. This indicates a more constant carbon supply for bacteria close to the roots during the growing season than in non-rhizosphere soil, including those groups that presumably prefer to utilize soil carbon.

In line with our second hypothesis, the stability of the net assimilation of plant-derived carbon in bacteria was not affected by the proximity to the roots. In contrast, saprophytic fungi showed a significantly lower stability of plant-derived carbon in non-rhizosphere soil than close to the roots. Nuccio et al. (2020) reported a strong difference in bacterial communities between rhizosphere and non-rhizosphere soil, while the fungal community differed more strongly between treatments with root carbon amendments. The relatively high temporal stability of plant-derived carbon in bacterial groups, being not different between rhizosphere and non-rhizosphere soil, might thus be explained by different bacterial communities; which are adapted to the different availability of plant-derived carbon over the season in both zones. In contrast to bacteria, the hyphal growth of saprophytic fungi would allow a similar portion of plant-derived carbon in rhizosphere and non-rhizosphere soil, comparable to AM fungi. Therefore, the strong differences in the net assimilation of plant carbon by fungi between rhizosphere and non-rhizosphere soils are likely attributable to different fungal communities, with the rhizosphere community taking advantage of constant carbon supply from roots, resulting in more stable net assimilation of plant carbon. Nuccio et al. (2020), further suggested that a functional succession of the rhizosphere community takes place, depending on the type and age of plant material provided. With regards to the bacterial community two aspects of our study support this hypothesis: 1) The relatively high temporal stability of bacterial groups compared to saprophytic fungi (Fig. 3), and 2) temporal stability being unaffected by the proximity to roots. A compositional community succession that enables the decomposition of different compound types of plant-derived carbon over the growing season would lead to a relatively constant net assimilation of plant-derived carbon, as observed in our study. Finally, this also supports the suggestion by Hannula et al. (2019) that seasonal shifts in bacterial communities are unlikely to change the functioning of the soils.

## Conclusions

Except AM fungi, being obligate symbionts with plants, all functional soil microbial groups were opportunistic in the energy sources (plant- and/or soil-derived) they used over the course of the growing season. Namely, the proportion of plant-derived carbon they assimilated, differs to some extent during the growing season. However, among different functional soil microbial groups, the ones which are more closely associated with the rhizosphere (in particular saprophytic fungi) showed higher variations in the net assimilation of plant-derived carbon, highlighting the importance of plant inputs for the soil functioning. Yet, irrespective of the variation in net assimilation of plant-derived carbon, the constant assimilation during the growing season suggests that succession in the microbial community might sustain soil biogeochemical cycling despite varying plant inputs, and we propose functional redundancy provisioned by succession by the microbial community as a mechanistic explanation.

## Supporting information

Supplementary Data und Figures

## Authors’ contributions

ML and GG conceived this study; MAR conducted the sampling and did the laboratory work. ML analyzed the data and wrote the manuscript. All authors contributed critically to the drafts and gave final approval for publication.

## Acknowledgment

We sincerely thank Agnes Fastnacht, Stefanie Hempel and Uta Gerighausen for maintaining and weeding the experimental site, Maria Wittwer and Steffen Rühlow for assistance with the GC-FID and GC-IRMS systems. We would also like to thank all former members of the Molecular Biogeochemistry research group who helped with the annual harvest of the experimental site. This study was financially supported by the Max Planck Institute for Biogeochemistry, Jena. Markus Lange gratefully acknowledges funding by the Zwillenberg-Tietz Foundation. Georg Dittmann, Dan Frederik Lange, Alice May Orme, Simon Andreas Schroeter and Carsten Simon would like to thank the International Max Planck Research School for Biogeochemical Cycles (IMPRS-gBGC) for funding. Simon Andreas Schroeter gratefully acknowledges the CRC 1076 AquaDiva, funded by the Deutsche Forschungsgemeinschaft (DFG).

