## Supplementary Data und Figures for "Uptake of plant-derived carbon and proximity to the root determine differences in temporal and spatial stability among microbial groups"

### Supplementary Information

Table S1: Simplified linear mixed effects models fitted with and without the factor “soil moisture” to assess how much is the temporal season effect (“time point”) driven by differences in soil moisture. Asterisks indicate the level of significance (“***” < 0.001, “**” < 0.01, “*” < 0.05, “.” < 0.1).

|  | LMM **without** soil moisture | LMM **with** soil moisture |
| --- | --- | --- |
|  | Chi^2^ | Chi^2^ |
| Actinobacteria | | |
| Soil moisture |  | 2.1 |
| Root | 2.1 | 3.3 . |
| Time point | 43.6 *** | 41.1 *** |
| G+ bacteria | | |
| Soil moisture |  | 1.2 |
| Root | 34.5 *** | 114.1 *** |
| Time point | 75.0 *** | 61.0 *** |
| G- bacteria | | |
| Soil moisture |  | 0.57 |
| Root | 49.18 *** | 54.46 *** |
| Time point | 24.28 *** | 18.52 *** |
| cy G- bacteria | | |
| Soil moisture |  | 1.5 |
| Root | 2.2 | 3.3 . |
| Time point | 13.1 ** | 12.9 ** |
| AM Fungi | | |
| Soil moisture |  | 0.5 |
| Root | 1.2 | 1.0 |
| Time point | 10.8 * | 12.6 ** |
| Saprophytic Fungi | | |
| Soil moisture |  | 3.5 . |
| Root | 16.2 *** | 20.3 *** |
| Time point | 10.7 * | 6.1 |
| Soil fauna | | |
| Soil moisture |  | 0.2 |
| Root | 4.8 * | 4.7 * |
| Time point | 31.3 *** | 31.4 *** |

Table S2: Average values and the respective standard deviation of individual lipid markers of concentration, d13C values and proportion of plant-derived carbon in plots differing in the vegetation type (C3 vs. C4). Also Pearson’s correlation coefficient of the individual markers to soil moisture is given, asterisks indicate the level of significance (“***” <0.001, “**”< 0.01, “*” < 0.05, “.” < 0.1).

|  |  | C3 Concentration [ µgC g^-1^] | | C4 Concentration [µgC g^-1^] | | C3 δ^13^C [‰] | | C4 δ^13^C [‰] | | Proportion of plant-derived carbon | | Correlation to soil moisture |
| --- | --- | --- | --- | --- | --- | --- | --- | --- | --- | --- | --- | --- |
| marker | group | rhizo | non-rhizo | rhizo | non-rhizo | rhizo | non-rhizo | rhizo | non-rhizo | rhizo | non-rhizo |  |
| **Me10C17** | actino | 1.67 ± 0.59 | 1.62 ± 0.51 | 1.85 ± 0.67 | 1.93 ± 0.60 | -28.6 ± 1.8 | -28.7 ± 1.2 | -22.2 ± 2.4 | -24.6 ± 2.8 | 0.44 ± 0.20 | 0.27 ± 0.16 | 0.49 *** |
| Me10C19 | actino | 1.00 ± 0.28 | 0.85 ± 0.23 | 1.04 ± 0.21 | 0.89 ± 0.27 | -28.7 ± 2.4 | -27.9 ± 1.4 | -21.6 ± 2.1 | -23.6 ± 2.9 | 0.48 ± 0.20 | 0.29 ± 0.19 | -0.05 |
| **C16;1w5** | neg | 1.64 ± 0.55 | 1.20 ± 0.42 | 2.04 ± 0.84 | 1.50 ± 0.51 | -25.2 ± 3.3 | -25.1 ± 1.6 | -16.8 ± 1.7 | -18.5 ± 2.1 | 0.58 ± 0.27 | 0.45 ± 0.14 | 0.35 ** |
| *C16;1w7* | neg | 2.20 ± 0.66 | 1.72 ± 0.59 | 2.73 ± 1.09 | 1.88 ± 0.56 | -28.1 ± 1.8 | -27.7 ± 1.4 | -20.0 ± 2.0 | -22.4 ± 2.6 | 0.55 ± 0.13 | 0.35 ± 0.17 | 0.23 . |
| **C17;1** | neg | 0.98 ± 0.38 | 0.91 ± 0.32 | 1.20 ± 0.46 | 1.09 ± 0.39 | -27.0 ± 2.8 | -26.8 ± 1.9 | -19.6 ± 3.0 | -22.1 ± 3.6 | 0.50 ± 0.22 | 0.32 ± 0.21 | 0.49 *** |
| *C18;1w7* | neg | 4.47 ± 1.56 | 3.15 ± 1.43 | 5.23 ± 2.22 | 3.59 ± 1.62 | -29.6 ± 1.3 | -29.1 ± 1.4 | -21.0 ± 1.8 | -24.0 ± 2.6 | 0.59 ± 0.13 | 0.34 ± 0.15 | 0.26 . |
| **C18;1w9** | neg | 3.69 ± 1.23 | 2.73 ± 1.02 | 4.20 ± 1.86 | 2.92 ± 0.95 | -28.5 ± 2.1 | -27.1 ± 1.7 | -19.4 ± 2.1 | -21.8 ± 3.0 | 0.62 ± 0.17 | 0.35 ± 0.13 | 0.36 ** |
| C17cy | neg (cy) | 0.99 ± 0.38 | 0.87 ± 0.28 | 1.22 ± 0.30 | 0.99 ± 0.26 | -26.9 ± 2.9 | -26.5 ± 1.9 | -18.5 ± 2.4 | -21.0 ± 3.1 | 0.57 ± 0.20 | 0.37 ± 0.21 | 0.13 |
| **C19cy** | neg (cy) | 1.99 ± 0.51 | 1.75 ± 0.65 | 2.20 ± 0.92 | 2.31 ± 0.56 | -30.4 ± 2.2 | -30.2 ± 1.3 | -23.2 ± 2.7 | -25.8 ± 2.7 | 0.49 ± 0.21 | 0.30 ± 0.16 | 0.28 * |
| C15a | pos | 1.82 ± 0.29 | 1.42 ± 0.36 | 1.99 ± 0.53 | 1.65 ± 0.37 | -27.1 ± 1.9 | -26.8 ± 1.5 | -19.0 ± 2.8 | -22.2 ± 3.0 | 0.55 ± 0.19 | 0.31 ± 0.17 | 0.19 |
| *C15i* | pos | 2.25 ± 0.63 | 1.94 ± 0.49 | 2.70 ± 0.66 | 2.31 ± 0.54 | -26.6 ± 2.1 | -26.2 ± 1.5 | -18.6 ± 2.7 | -21.2 ± 3.4 | 0.54 ± 0.16 | 0.34 ± 0.20 | 0.24 . |
| C16i | pos | 1.17 ± 0.33 | 0.91 ± 0.25 | 1.28 ± 0.27 | 1.01 ± 0.25 | -26.9 ± 2.5 | -26.3 ± 1.8 | -18.8 ± 2.9 | -21.0 ± 3.6 | 0.55 ± 0.20 | 0.36 ± 0.19 | 0.08 |
| **C17a** | pos | 0.77 ± 0.25 | 0.64 ± 0.21 | 0.88 ± 0.27 | 0.73 ± 0.22 | -24.4 ± 2.8 | -24.2 ± 2.0 | -16.4 ± 3.0 | -19.2 ± 3.6 | 0.54 ± 0.22 | 0.34 ± 0.21 | 0.29 * |
| **C17i** | pos | 0.74 ± 0.23 | 0.68 ± 0.21 | 0.92 ± 0.22 | 0.79 ± 0.22 | -25.2 ± 3.3 | -25.2 ± 2.6 | -17.8 ± 3.7 | -20.4 ± 4.2 | 0.50 ± 0.22 | 0.33 ± 0.21 | 0.31 * |
| **C18;2w6** | fungi | 1.70 ± 0.69 | 0.99 ± 0.79 | 2.22 ± 1.39 | 1.03 ± 0.47 | -29.7 ± 3.4 | -28.7 ± 4.1 | -19.2 ± 1.9 | -21.9 ± 4.5 | 0.72 ± 0.21 | 0.47 ± 0.23 | 0.29 * |
| C16;1w5NL | AM Fungi | 3.46 ± 2.76 | 3.26 ± 2.58 | 4.70 ± 5.22 | 3.23 ± 3.13 | -30.5 ± 1.2 | -30.4 ± 1.2 | -18.6 ± 1.8 | -18.7 ± 1.8 | 0.81 ± 0.10 | 0.80 ± 0.09 | -0.05 |
| C20;4w6 | fauna | 0.39 ± 0.11 | 0.35 ± 0.12 | 0.51 ± 0.16 | 0.38 ± 0.12 | -29.5 ± 4.6 | -29.6 ± 3.4 | -21.1 ± 5.1 | -25.3 ± 4.9 | 0.57 ± 0.35 | 0.29 ± 0.28 | -0.05 |

**Table S3:** Average stability and S.E. of the temporal stability of plant-derived carbon assimilated and the microbial biomass. Numbers correspond to Fig 3.

| Microbial group | Proximity to Roots | Concentrations | | Fp values | |
| --- | --- | --- | --- | --- | --- |
|  |  | Average | Average | S.E. | S.E. |
| Actinobacteria | non-rhizosphere | 4.61 | 4.27 | 1.21 | 0.52 |
|  | rhiz | 6.02 | 4.37 | 1.01 | 0.84 |
| Soil Fauna | non-rhizosphere | 4.40 | 2.13 | 1.00 | 1.65 |
|  | rhizosphere | 5.86 | 6.70 | 5.50 | 1.57 |
| cy G- bacteria | non-rhizosphere | 4.75 | 3.31 | 0.87 | 0.55 |
|  | rhizosphere | 4.76 | 3.66 | 0.77 | 0.60 |
| G- bacteria | non-rhizosphere | 4.03 | 7.38 | 2.22 | 0.26 |
|  | rhizosphere | 4.97 | 5.36 | 0.77 | 0.38 |
| G+ bacteria | non-rhizosphere | 4.66 | 4.44 | 1.00 | 0.31 |
|  | rhizosphere | 5.59 | 6.44 | 1.54 | 0.41 |
| Saproph. Fungi | non-rhizosphere | 2.00 | 2.04 | 0.39 | 0.28 |
|  | rhizosphere | 2.72 | 3.83 | 0.72 | 0.27 |
| AM fungi | non-rhizosphere | 2.19 | 11.85 | 1.25 | 0.56 |
|  | rhizosphere | 1.58 | 9.29 | 1.34 | 0.25 |

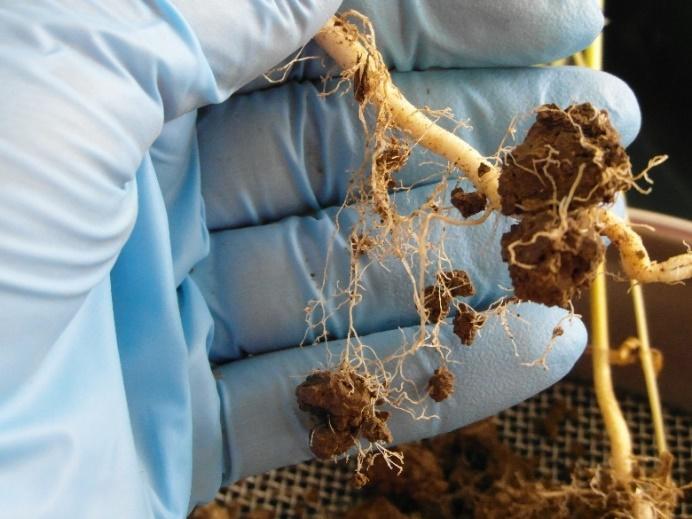

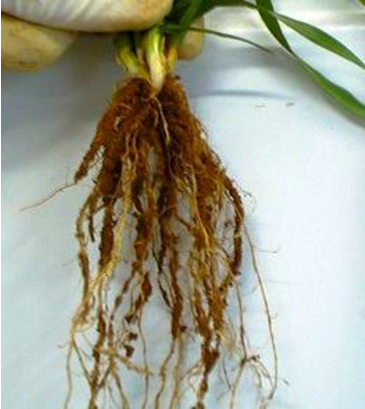

**Figure S1:** Photographic depicts the sampling of rhizosphere soil. All soil that was attached to the roots was considered as rhizosphere soil.

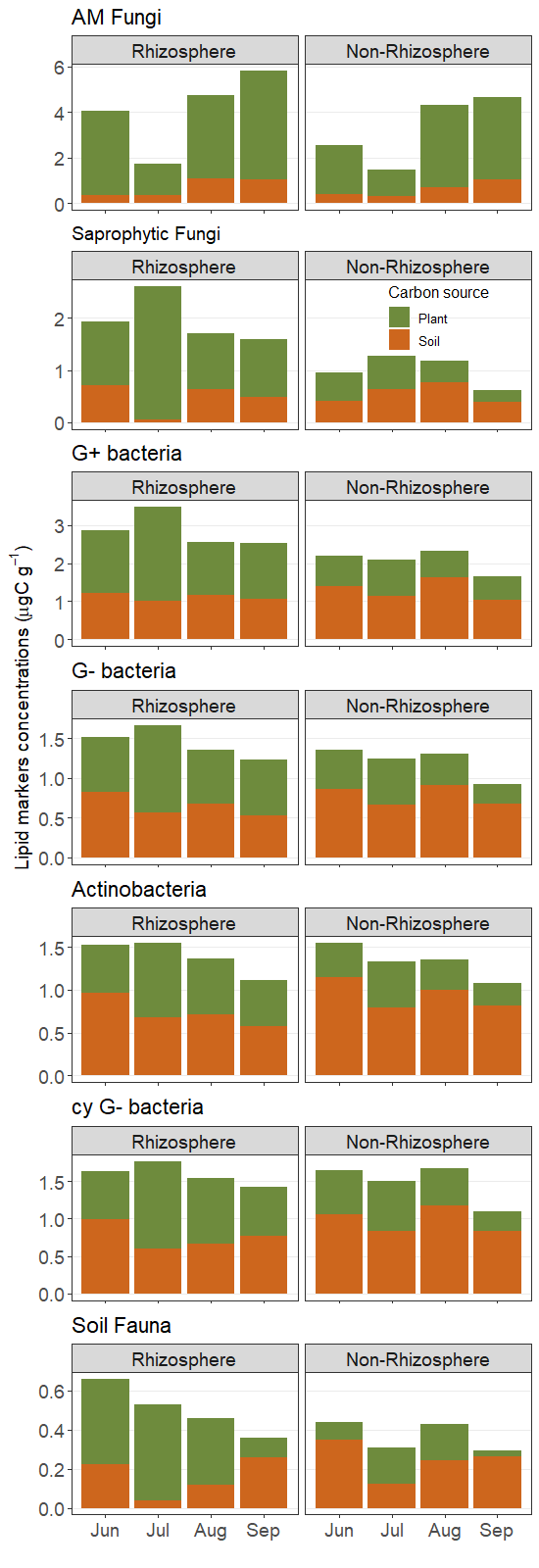

**Figure S2:** Biomass of microbial groups over the growing season in rhizosphere and non-rhizosphere soil and the contribution of plant and soil derived carbon. The total biomass corresponds to the values in Fig. 1, the portion is calculated based on the Fp values presented in Fig. 2.
